# MetaDAG: a web tool to generate and analyse metabolic networks

**DOI:** 10.1101/2024.05.15.593827

**Authors:** Pere Palmer-Rodríguez, Ricardo Alberich, Mariana Reyes-Prieto, José A. Castro, Mercè Llabrés

## Abstract

We introduce MetaDAG, a web-based tool designed for metabolic network reconstruction and analysis. MetaDAG is capable of constructing metabolic networks associated with specific organisms, sets of organisms, sets of reactions, sets of enzymes, and sets of KO (KEGG Orthology) identifiers. To generate these metabolic networks, MetaDAG retrieves from the KEGG database the chemical reaction information that corresponds to the user’s queries. MetaDAG computes a reaction graph as a first metabolic graph model. This reaction graph is a network in which nodes represent reactions, and edges between reactions indicate the presence of a metabolite produced by one reaction and consumed by another. Next, as a second metabolic model, MetaDAG computes a directed acyclic graph called a metabolic DAG (m-DAG for short). The m-DAG is obtained from the reaction graph by collapsing all strongly connected components into single nodes. As a result, the m-DAG representation reduces considerably the number of nodes while keeping and also highlighting the network’s connectivity. Both metabolic models, the reaction graph, and the m-DAGs, are displayed on an interactive web page to assist the users in visualising and analysing the networks. Furthermore, users can retrieve the node’s information linked to the KEGG database. All generated files, including images containing metabolic network information and analysis results, are available for download directly from the web page. In the Eukariotes test presented here, MetaDAG has demonstrated its effectiveness in classifying all eukaryotes from the KEGG database at both the kingdom and phyla taxonomy levels.

## Introduction

In the current omics era, the focus of metabolic function analysis has shifted from individual organisms to entire microbial communities, as well as the intricate metabolic interactions among community members. The vast quantity of sequence data generated from metagenomic and metatranscriptomic samples requires specialized tools for the taxonomic and functional annotation of genomes, genes, and proteins, as well as their subsequent integration and analysis. In this context, the reconstruction and analysis of metabolic networks are crucial for understanding the metabolic profiles and their interactions within microbial communities. These networks can be extensive, often encompassing hundreds or even thousands of interrelated reactions. As a result, the development of bioinformatic tools for their visualization and comprehensive analysis has become a necessity.

Several methodologies for metabolic reconstruction have been developed, including machine learning-based approaches that predict and reconstruct metabolic pathways (see [1] and [2] for an extensive review of these methods). Nevertheless, the constantly increasing amount of metabolic information stored and characterized in several public repositories according to their functions, such as KEGG [3], MetaCyc [4], BioCyc [5], among others, strengthens the need for automated metabolic reconstruction methods based on curated metabolic data information. In this line of research, various approaches to metabolic network reconstruction, analysis, and comparison can be found in the literature [6], [7](see [8–10] for surveys on different approaches and tools). Each approach selects a representation of metabolic networks that models information of interest, proposes a similarity or a distance measurement, and possibly supplies a tool. Models of metabolic networks range from a high-level abstraction of metabolic information, such as abstract metabolic networks, to a lower level of abstraction, such as reaction networks, metabolic hypergraphs, or stoichiometric matrices.

In [11] a novel methodology for the analysis of metabolic networks was presented. This methodology incorporates the concept of strongly connected components in a reaction graph, called *metabolic building blocks*. Additionally, a new model of metabolic networks, known as a *metabolic DAG*, was defined to combine the reaction graph information with the network’s topology.

This methodology has been used in different contexts, and proven to be successful in comparing, analysing, and visualising metabolic networks, as shown in previous studies [12,13]. As a result, we present here MetaDAG, a web-server tool that implements the metabolic DAG methodology. MetaDAG automates the metabolic network reconstruction using several different identifiers of data stored in the KEGG database, which we choose as our source of metabolic information due to its curated nature and standardized presentation.

MetaDAG offers metabolic network reconstructions from various queries, including specifying a single organism, a group of organisms, a set of reactions, a set of enzymes, or a set of KO (KEGG Orthology) identifiers. This capability encompasses the reconstruction of metabolic networks from single microbial samples to consortia of two or more organisms, and even to complex metagenomic samples, providing a versatile tool for different analytical scenarios.

Metabolic networks are generated by retrieving the reactions associated with user-specified queries from the KEGG database. Initially, it computes a metabolic network as a reaction graph. From the reaction graph, a directed acyclic graph called a metabolic DAG (m-DAG for short) is computed by collapsing all strongly connected components of the reaction graph into single nodes, called a metabolic building block (MBB for short). In the m-DAG, two MBBs are connected through an edge if there is at least one pair of reactions (one in each MBB) connected by an edge in the reaction graph. As a result, the m-DAG representation reduces considerably the number of nodes while keeping the network’s connectivity. Hence, at first glance, this m-DAG representation offers an easy-to-interpret topological analysis of the reconstructed metabolic network. Both models of metabolic networks, the reaction graphs and the m-DAGs, are displayed in an interactive web to aid users in visualising and analysing the networks, as well as to retrieve the node’s information linked to KEGG.

Furthermore, when examining different groups of organisms, experiments, or samples, MetaDAG also calculates the core and pan metabolism associated with each group. It then provides the results of a comparative analysis of their respective m-DAGs. This comparative analysis offers valuable insights into the shared and unique metabolic features across different groups, experiments, or samples, as shown in the results of the Eukaryotes test, conducted to assess the tool’s performance and functionality. Given the vast amount of information MetaDAG generates for each query, a comprehensive user guide in R is provided to assist users in understanding, handling, and analysing all the results.

## Design and implementation

In this section, we first recall the methodology of metabolic networks and the construction of metabolic DAGs, and we present its implementation in the proposed tool *MetaDAG* to generate, analyse, and compare metabolic networks.

### Reaction graphs & Metabolic DAG models

A *reaction graph model* of metabolism, is a directed graph *G*_*R*_ = (*R, E*) in which its set of nodes is the set *R* of chemical reactions present in the metabolism, and its set of edges, *E*, is defined as follows: there is an edge from *R*_*i*_ to *R*_*j*_ if, and only if, there is at least one metabolite produced by *R*_*i*_ that is consumed by *R*_*j*_. Reversible reactions are modeled by two different nodes, one for the forward reaction and the other for the backward reaction. When a directed graph has no cycles, it is called a *directed acyclic graph*, DAG for short.

A *path* from a node *u* to a node *v* in a directed graph *G* is a sequence of nodes {*u*_0_, *u*_1_, …*u*_*k*_} such that *u*_0_ = *u, u*_*k*_ = *v* and (*u*_*i*_, *u*_*i*+1_) is an edge in *G* for *i* = 0, …, *k*− 1. Two nodes *u, v* are said to be biconnected if there is a path in each direction between them. A *strongly connected component* of a directed graph *G* is a subgraph such that every pair of nodes in it are biconnected, and it is maximal under inclusion with this property [14,15]. Since biconnectivity is an equivalence relation, the collection of strongly connected components forms a partition of the set of nodes of *G*. If each strongly connected component is contracted to a single node, the resulting quotient graph is a DAG, the *condensation* of G. Notice that there is an arc from a node *s*_*i*_ to a node *s*_*j*_ in the condensation of a directed graph *G* if, and only if, there is an arc in *G* from a node *u* ∈ *s*_*i*_ to a node *v* ∈ *s*_*j*_. Thus, for every reaction graph *G*_*R*_, we can consider its collection of strongly connected components and compute its condensation, which will become a DAG. We call a *metabolic building block* (MBB for short) each strongly connected component in the reaction graph *G*_*R*_, and we call *metabolic DAG* (m-DAG for short) the condensation of a reaction graph *G*_*R*_. *Essential MBBs* are the MBBs with only one reaction where its deletion increases the number of connected components of the m-DAG.

Given a set of reaction graphs, its *pan-metabolism* is the reaction graph obtained when considering the reactions that belong to one or more reaction graphs in the set. The *core-metabolism* is the reaction graph obtained when considering the reactions that belong to all reaction graphs in the set. Intuitively, the pan-metabolism represents the joint metabolism while the core-metabolism represents the shared metabolism. These terms were inspired by the pangenome concept [19].

Figure 1 shows the reaction graph of the glycolysis pathway in *Homo sapiens*, computed with the tool metaDAG. Notice that we show in this figure the reaction graph of a single pathway, which is a very reduced set of reactions compared to the complete metabolism of *Homo sapiens* (35 reactions versus 1860). The metabolic DAG associated with this reaction graph is shown in Figure 2. This m-DAG has 7 MBBs, 3 MBBs consisting of only one reaction, with 2 of them being essential MBBs, and 4 MBBs with more than one reaction. To easily explore the reactions that belong to an MBB, users can select it, and a small window on the right presents information for all the reactions within the MBB. This includes the KEEG ID of the reaction, its chemical formula, and a hyperlink pointing to the reaction information on the KEGG webpage.

**Figure 1:**
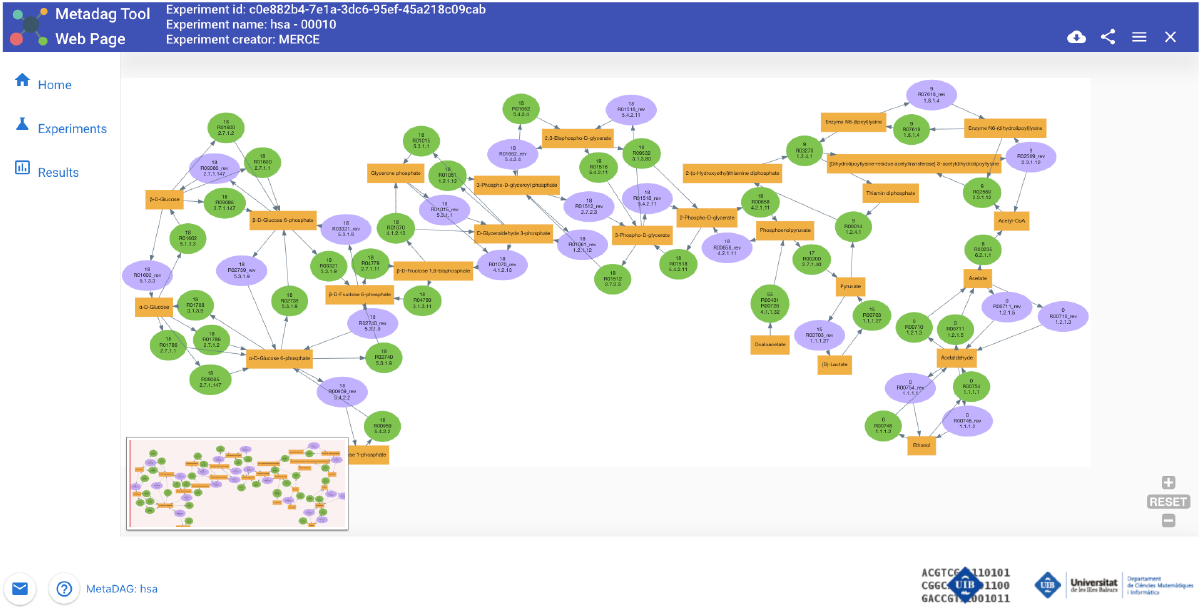
Reaction graph of the glycolysis pathway in Homo sapiens, computed and displayed with the tool MetaDAG.

**Figure 2:**
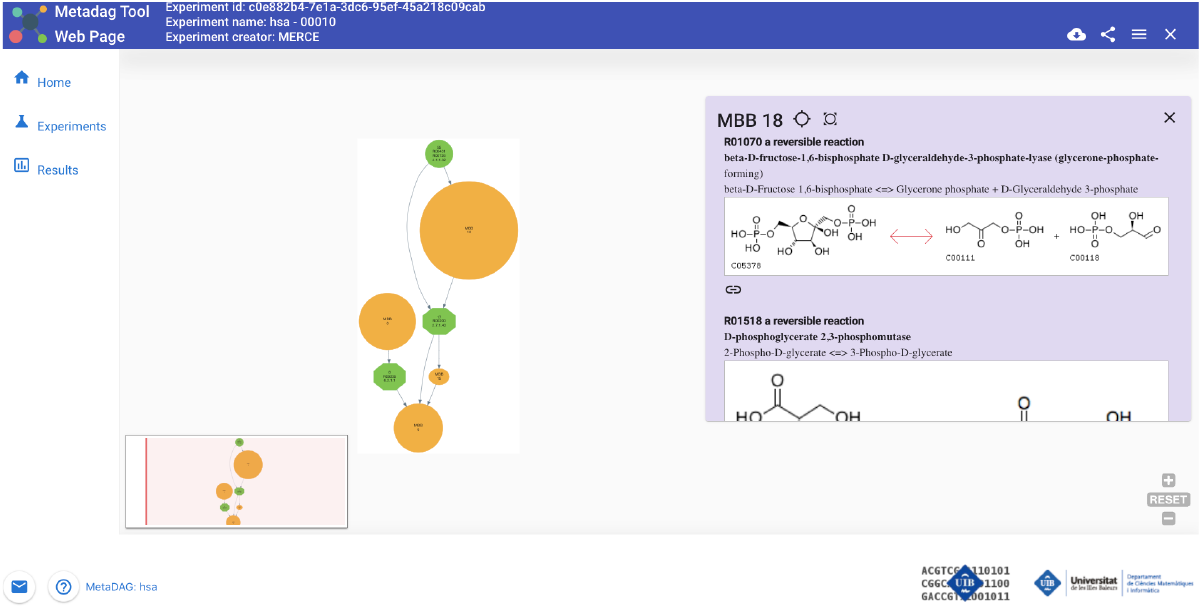
Metabolic DAG corresponding to the reaction graph of the glycolysis pathway in *Homo sapiens*

### MetaDAG’s calculation and outcome scope

MetaDAG provides the reaction graphs and the corresponding metabolic DAG models under the following queries:

#### 1. A specific pathway of an organism

To generate a reaction graph and m-DAG, users are required to choose one pathway and an organism from the list of metabolic pathways and organisms available in the KEGG database. Then, all reactions present in the selected pathway for the chosen organism are taken into account to construct the reaction graph and m-DAG.

#### 2. Global metabolic network of one organism

This query generates the reaction graph and m-DAG for the entire metabolism of a single organism. In this query, the focus is on selecting a singular organism, and the metabolic models are constructed by incorporating all reactions present across all pathways of the chosen organism.

#### 3. Specific pathway of all organisms

In this query, a single pathway from the KEGG catalog is chosen. The metabolic models are then constructed by considering all reactions present in that pathway for at least one organism. Essentially, this query provides the reaction graph and m-DAG for a reference pathway sourced from KEGG.

#### 4. Global metabolic network of a list of organisms

The user has to provide a list of organisms using their KEGG identifier from the KEGG database. Then, the reaction graph and m-DAG of each organism are constructed in the same manner as in the second experiment. Additionally, the pan-metabolism and core-metabolism of all organisms in the provided list are generated. Users also have the flexibility to select specific groups of organisms from the list, and for each group, their pan and core metabolisms are constructed as well. Furthermore, users can request similarities among all m-DAGs as explained in the next section.

#### 5. Synthetic metabolism of a list of compounds, reactions, enzymes, or KOs

This query generates the reaction graph and m-DAG for a given set of compounds, reactions, enzymes, or KO identifiers. When a set of enzymes is provided through the EC number or a set of KO identifiers, or when a list of reactions is specified, even if they are not necessarily from the same organism, metaDAG constructs the reaction graph by retrieving all relevant information from KEGG. This includes enzymes, reactions, and compounds associated with the input data.

Similarly, when a list of compounds is provided, the models are constructed by considering all reactions where a compound from the list is present in either its product or substrate. Additionally, users can request the similarities among all m-DAGs.

#### 6. Comparison of several experiments

This query allows joining the previous queries. Namely, the user may compare a set of organisms and a set of reactions, enzymes, KOs, or compounds and obtain the analysis of the whole experiment. That is the pan and core metabolisms. In addition, the metabolisms of a synthetic organism can also be constructed. Finally, users can request the similarities among all m-DAGs.

### Pairwise similarity of m-DAGs

For every pair of m-DAGs, the tool provides two similarity measures based on the similarity of their MBBs: the *Munkres-similarity* and the *MSA-similarity*.

The Munkres-similarity is the similarity defined in [11] which we briefly recall here. Firstly, to define a similarity score of two MBBs, the pairwise reactions’ similarity score as it was defined in [16] is considered. Next, given two MBBs, *MBB*_1_ and *MBB*_2_ its similarity score, *S*_*mbb*_(*MBB*_1_, *MBB*_2_), is computed by:

- defining a complete bipartite graph in which the reactions in *MBB*_1_ and *MBB*_2_ are nodes and the weight of each edge (*R*_*i*_, *R*_*j*_) ∈ *MBB*_1_ × *MBB*_2_ is the similarity of *R*_*i*_ and *R*_*j*_;
- applying the maximum weighted bipartite matching algorithm to the resulting graph to obtain the best match between *MBB*_1_ and *MBB*_2_;
- summing the scores of the best match and dividing it by max{|*MBB*_1_|, |*MBB*_2_|}.

Finally, the similarity measure between two m-DAGs, *Sim*(*mD*_1_, *mD*_2_) is computed by:

- defining a complete bipartite graph in which the MBBs in *mD*_1_ and *mD*_2_ are nodes and the weight of each edge (*MBB*_*i*_, *MBB*_*j*_) ∈ *mD*_1_ × *mD*_2_ is *S*_*mbb*_(*MBB*_1_, *MBB*_2_);
- applying the maximum weighted bipartite matching algorithm to the resulting graph to obtain the best match between *mD*_1_ and *mD*_2_;
- summing the scores of the best match and dividing it by max{|*mD*_1_|, |*mD*_2_|}.

The MSA-similarity, which stands for Maximum Similarity Assignment, is defined as follows: Let *MBB*_1_ and *MBB*_2_ be two MBBs, its similarity score is

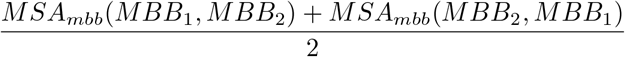

where *MSA*_*mbb*_(*MBB*_1_, *MBB*_2_) is defined by

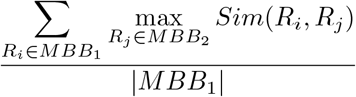

and, analogously, we define *MSA*_*mbb*_(*MBB*_2_, *MBB*_1_). Again, the pairwise reactions’ similarity score is the similarity defined in [16]. Then, the similarity measure between two m-DAGs, *mD*_1_ and *mD*_2_ is

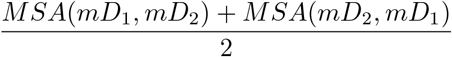

where *MSA*(*mD*_1_, *mD*_2_) is defined by

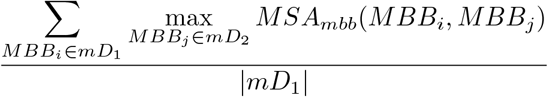

and, analogously, we define *MSA*(*mD*_2_, *mD*_1_).

We discuss and compare these two similarities in the results section.

### MetaDAG output

For every query, MetaDAG provides the corresponding metabolic models, that is, the reaction graphs and the m-DAGs. In addition, MetaDAG provides the pan and the core reaction graphs and m-DAGs of different experiments or organisms, as well as the pairwise similarities of all m-DAGs. Upon completion of a query, users receive an email containing a job ID, which they can use to access the results on the MetaDAG front page. In the upper-right panel, users have the option to download the results, share them, or display them through user-friendly icons. By clicking the display icon, a new window opens, allowing users to select any computed metabolic graph for a specific organism or experiment. This includes the m-DAG, reaction graph, and the largest connected component of each m-DAG. Once a metabolic graph is chosen, it is displayed on an interactive webpage to facilitate user exploration and analyses. For detailed instructions on interpreting the results of the MetaDAG tool, please refer to the pipeline available at https://github.com/biocom-uib/MetaDag/tree/main.

### Metabolic graphs displayed on the webpage

#### Reaction Graph

To help users to visualise the reaction graph and contextualise the reactions in it, metaDAG shows not only the reactions, i.e., the nodes in the reaction graph, but also the metabolites associated with each reaction in the KEGG database. Reactions are depicted as green circular nodes, and, for every reversible reaction, a new node depicted by a purple oval is added. The directed edges from a metabolite to a reaction, or vice versa, show if the metabolite is a substrate or a product of the corresponding reaction. Every reaction and compound can be selected, and then, a new window emerges with its information as well as a link pointing to the corresponding information on the KEGG’s webpage.

#### m-DAG

The nodes of the m-DAGs, i.e., the metabolic building blocks (MBBs), are displayed with different types of nodes. Essential MBBs are depicted as green octagons, while green circles are the MBBs with only one reaction. Yellow circles are those MBBs with more than one reaction, and their size refers to the number of reactions in the MBB. To easily know the reactions that belong to an MBB, the user can select it and a small window appears with the information of all reactions in the MBB. Namely, the ID of the reaction in KEGG, its chemical formula, and a link pointing to the reaction information in the KEGG webpage. On the top-right of this window, in the third icon, the user may contextualize the reactions, since these selected reactions appear highlighted over the reference pathway in the KEGG format.

#### Biggest Connected Component

When an m-DAG is huge, as in the case of the m-DAG of the entire metabolism of one organism, there are often many isolated nodes. However, as discussed in the results section, these m-DAGs have a considerably large connected component. Therefore, we found it valuable to separately extract the biggest connected component of an m-DAG, which is displayed in the same format as the full m-DAG. In the supplementary material, File S1 presents the core m-DAG for the kingdom of Animalia, whereas File S2 displays its largest connected component.

### Downloadable metaDAG data results

All results for a given query can be classified into two categories: On one side, some files contain information about the relationships between the m-DAGs, the MBBs, and the various organisms or samples provided. On the other side, several files describe each m-DAG, so that the information can be visually represented. MetaDAG provides a specific viewer to display the m-DAGs, and also the Reaction Graphs, giving users the possibility to carefully analyze those graphs. Additionally, there is an option to download the files containing the data. In this case, users can select which items they wish to download.

In the most extensive case, the available information is organized into the following categories:

- General data: Files with the information describing the whole set of m-DAGs, they contain: for each m-DAG which MBB it contains, for each MBB which are the reactions involved, and, to simplify the data handling, a third file having a combination of both views, that is: for each m-DAG which reactions it has and in which MBB.
- Representation data: A detailed representation of each generated metabolic network (m-DAG and reaction graph), including the different format representations of the graphs, such as, graphml, svg, csv, etc.

We refer to File S3 in the supplementary material for a full description of every item that can be downloaded.

### Implementation

MetaDAG is an innovative web-based tool designed to provide users with a powerful and efficient way to analyse and visualise complex DAGs modelling a metabolic network. The client-side, or front-end, of MetaDAG has been crafted using Angular and TypeScript, ensuring a robust and responsive user interface. The Angular framework enables consistent integration of components, bringing a dynamic and interactive user experience. TypeScript, a superset of JavaScript, further enhances the front-end development process promoting better code organization, early error detection, and improved maintainability.

On the server-side, the back-end of MetaDAG has been engineered using Java, a versatile and highly reliable programming language, in combination with the popular Spring framework. The Spring framework simplifies the development process by providing a comprehensive set of tools and libraries for building scalable and performant applications. With Spring, the back-end of MetaDAG benefits from features such as dependency injection and aspect-oriented programming, making it a solid foundation for data-intensive operations.

The back-end has been designed in two layers. The front layer is responsible for serving data requests from the front-end (e.g.: serving the files requested o preparing the data the user wants to download). Furthermore, when a user makes a new query, the front layer prepares the data needed to fulfill the request and passes them to the calculation layer, wherein an independent process of the calculations is performed. This way, the front layer can serve customer requests quickly. The calculations are made independently and they can last a significant amount of time, so a contact email for the user has to be provided. When the calculations are finished an email giving access to the results is sent.

MetaDAG makes use of the fundamental biological information available on the Kyoto Encyclopedia of Genes and Genomes (KEGG) database. KEGG is a widely recognized and highly curated database that contains an extensive collection of biological pathways and networks. It provides standardized nomenclature and annotations for genes, proteins, enzymes, orthologs, and pathways. We consider the KEGG database since it is an extensive, reliable, and widely used resource explicitly designed to present data in a standardized way.

## Results

The MetaDAG’s methodology has been already applied in two different scenarios. First, MetaDAG was successfully applied to obtain the metabolic DAGs of 2,328 symbiotic genomes included in the public database *Symbiotic Genomes Database - SymGenDB* [12] available at http://symbiogenomesdb.uv.es/. All metabolic DAGs as well as the corresponding pan and core metabolisms at the genus level were calculated and stored in the Meta-DAGs section of the database.

Second, to explore the tools’ usability regarding the metabolic network’s topology, in [13] the reaction graph and the m-DAG of the minimal metabolic network were constructed from the theoretical minimal gene set machinery revised in [17]. The reaction graph of this minimal metabolic network consisted of 80 compounds and 98 reactions, while its m-DAG had 36 MBBs. Additionally, 12 essential reactions were identified in the m-DAG that are critical for maintaining the connectivity of the network. Similarly, the m-DAG of JCVI-syn3.0, and of Candidatus *Nasuia deltocephalinicola* were constructed. JCVI-syn3.0 is an artificially designed and manufactured viable cell whose genome arose by minimizing the one from *Mycoplasma mycoides* JCVI-syn1.0 [18], and Candidatus *Nasuia deltocephalinicola* is the bacteria with the smallest natural genome known to date. The comparison of the m-DAGs derived from a theoretical, an artificial, and a naturaly reduced genome denoted slightly different lifestyles, with a consistent core metabolism.

### Eukaryotes test

To further evaluate our tool’s usability, as well as the m-DAG methodology, we ran a test considering all Eukaryotes from the KEGG database. We introduced them as a list of organisms using query number 4th on the metaDAG frontpage. Currently, Eukaryotes are distributed in 535 Animals, 154 Fungi, 139 Plants, and 56 Protists (see File S4 in the supplementary material). Hence, we obtained the reaction graphs and the m-DAGs of every organism, as well as the pan and core reaction graphs and m-DAGs for every Kingdom group. In addition, the Munkres-similarity and the MSA-similarity for every pair of m-DAGs were calculated. It took 244 minutes to obtain the results from the tool running on an AMD/7282 biprocessor provided with 512 GiB RAM.

Interestingly, we found that the core reaction graph of all Eukaryotes is empty, which means the absence of any reaction shared across all Eukaryotic organisms. However, the core reaction graphs within the kingdom taxonomy level were not empty. Table 1 shows the number of reactions shared between the kingdoms. We can observe that Animals share 133 reactions and 117 MBBs, Plants share 303 reactions and 249 MBBs, while Fungi and Protists both share 25 reactions and 22 and 21 MBBs, respectively.

**Table 1:**
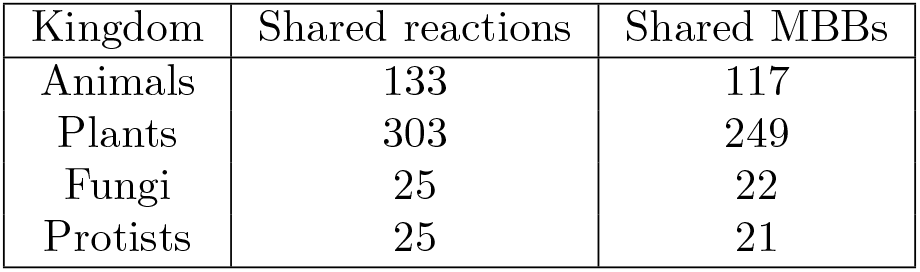
This table shows the number of reactions shared by all organisms in every kingdom taxonomy level as well as the number of shared MBBs. This corresponds to the number of nodes in the core reaction graph and core m-DAG for every kingdom group.

| Kingdom | Shared reactions | Shared MBBs |
| --- | --- | --- |
| Animals | 133 | 117 |
| Plants | 303 | 249 |
| Fungi | 25 | 22 |
| Protists | 25 | 21 |

Regarding the topology of the m-DAGs, we observe that all computed m-DAGs strongly share a topology profile. Namely, they have many isolated nodes, most of them consisting of MBBs with only one reaction, and also, they all have a considerably big connected component. Figure 3 on the left, reflects this profile. On the x-axis, we ordered the connected components of every m-DAG from the biggest to the smallest one regarding its node size. On the y-axis, we show the number of nodes of every connected component. We show the results for every kingdom in a colour palette: Animals in red lines, Plants in green, Fungi in yellow, and Protists in black. We can observe that most lines are plotted together, meaning that they have the same or quite a similar number of nodes per connected component. The animals have the biggest component, with 640 nodes, followed by the plants, with 597 nodes. We can also observe, that all m-DAGs rapidly descend to connected components with only one node. Again, the animals have the biggest number of connected components, but most of them have only one node. To enhance the visualization of connected components with more than one node, we adjust the results by rescaling them using a logarithmic scale. Figure 3 on the right shows the results. We observe that Animals have the biggest connected components, followed by Plants, Fungi, and Protists. We refer to the pipeline available at *https://github.com/biocom-uib/MetaDag/tree/main* for a complete analysis of these graph’s topology.

**Figure 3:**
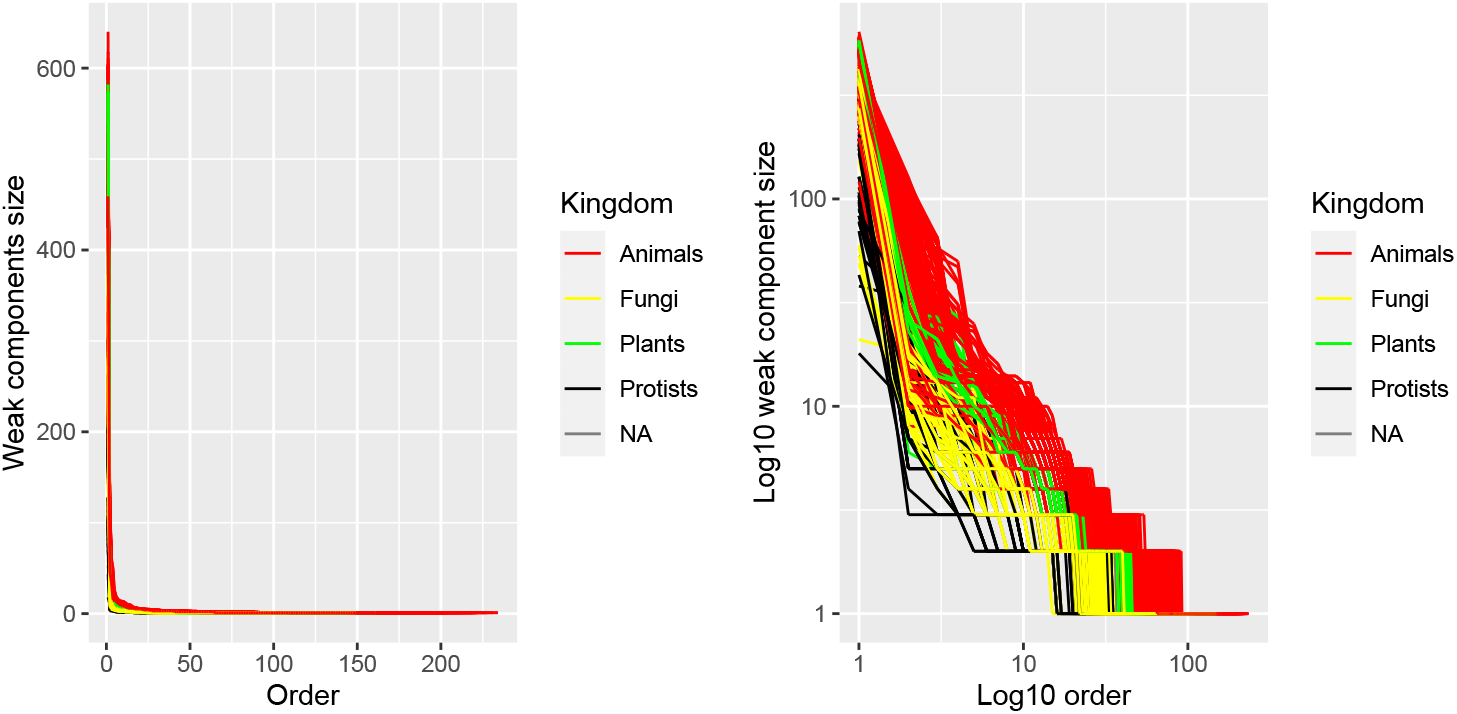
Size of the connected components of every m-DAG (y-axis) ordered in decreasing order (x-axis).

Concerning the number of reactions in each node of every m-DAG, we also obtain the same pattern as before. Specifically, in all kingdoms, all m-DAGs have a huge node (MBB) with 979 reactions. And, the majority of the MBBs have only one or two reactions. For further details, refer to File S5 in the Supplementary Material.

As for the large-scale comparison of all constructed m-DAGs, both, the MSA-similarity and Munkres-similarity measures were calculated with MetaDAG. In Figure 4 we show the corresponding heatmaps to visualize the results. A heatmap allows for a visual rendering of the similarity matrix produced by each similarity measure. Each cell (*i, j*) in the heatmap shows the similarity value between the *i*-th and *j*-th m-DAGs, colour-coded so that darker colours correspond to a high degree of similarity (from dark blue to yellow). In our case, the heatmaps are symmetric and cells in the main diagonal always show the darkest colour, resulting from comparing an organism with itself. We can observe in this figure that, both similarity measures correctly classify m-DAGs at the kingdom level, and we also clearly distinguish two separate groups within the Animals kingdom. Hence, for each similarity measure, we computed the hierarchical clustering of all Eukaryotes, using the Ward method with 4 clusters. The results are shown in Table 2. We observe that the MSA-similarity separates almost all Animals (except 9 out of 535) into two homogeneous and distinct clusters. Plants are also separated (except 14 from 139) in another homogeneous and distinct cluster, while Fungi and Protists end up within the same cluster together with the 9 animals and 14 plants. These 9 animals are nematodes or flatworms and exhibit a parasitic nature, leading to the development of various diseases. This parasitic condition could result in significant differences in their metabolic characteristics compared to other organisms. Concerning the 14 plants, they are all the green and red algae in the KEGG database. Hence, they are all clustered together. Some of them contain high lipid content and are suitable for biodiesel production. As in the animal’s case, all of them exhibit unique characteristics that can influence their metabolism compared to other plants.

**Table 2:** Clusters obtained at the kingdom level for all m-DAGs of Eukariotes with the MSA and Munkres-similarity measures.

| MSA-similarity |  |  |  |  |
| --- | --- | --- | --- | --- |
| Clusters | Animals | Fungi | Plants | Protists |
| 1 | 331 | 0 | 0 | 0 |
| 2 | 195 | 0 | 0 | 0 |
| 3 | 0 | 0 | 125 | 0 |
| 4 | 9 | 154 | 14 | 56 |

| Munkres-Similarity |  |  |  |  |
| --- | --- | --- | --- | --- |
| Clusters | Animals | Fungi | Plants | Protists |
| 1 | 331 | 0 | 0 | 0 |
| 2 | 197 | 0 | 0 | 0 |
| 3 | 0 | 0 | 125 | 0 |
| 4 | 7 | 154 | 14 | 56 |

**Figure 4:**
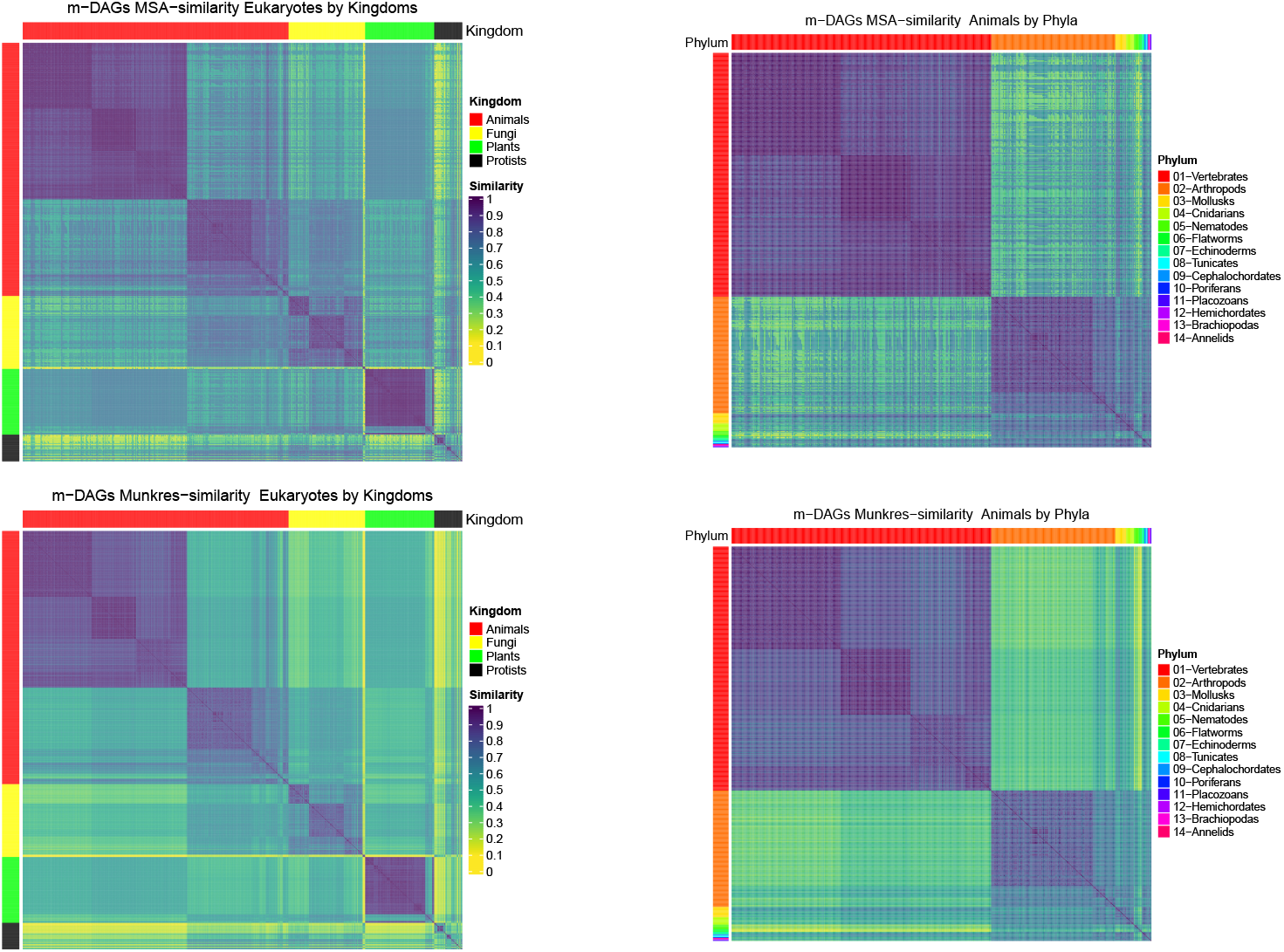
Heatmaps of the similarity matrices of KEGG Eukaryotes: (a) MSA-similarity at the kingdom level (top-right); (b) Munkres-similarity at the kingdom level (bottom-right); (c) MSA-similarity of animals at the Phylum level (top-left); (d) Munkres-similarity of animals at the Phylum level (bottom-left)

Similar results are also obtained with the Munkres-similarity. Indeed, as shown also in Table 2, in this case, we obtain only 7 of the previous 9 animals clustered together with all Fungi, Protists, and green and red algae. See Files S6 and S7 in the Supplementary material for a detailed description of these organisms classification.

From this classification, a pertinent question arises: What factors cause these algae and animals to be distinguished from their respective kingdoms? To address this query, we revisited the core metabolism obtained with MetaDAG, but this time for each cluster rather than the kingdom’s core metabolism. Table 3 displays the number of reactions and MBBs shared within each cluster.

**Table 3:**
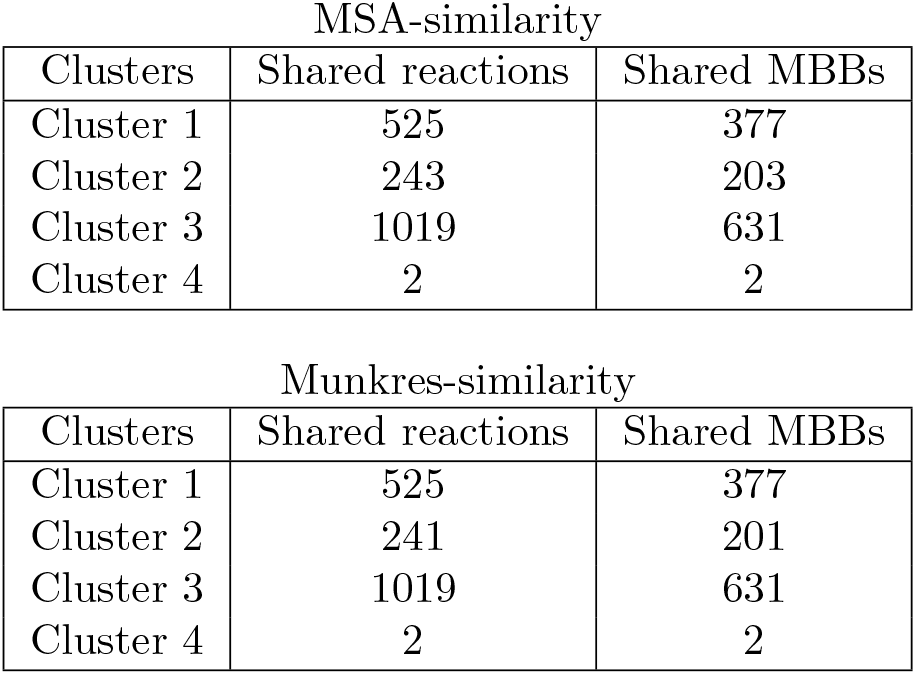
Reactions and MBBs shared by clusters.

We now observe that by dividing animals into two clusters - vertebrates and invertebrates - the number of shared reactions increases substantially. Notably, the vertebrates cluster shares 377 MBBs, while the number of shared reactions is 525. This indicates that vertebrates share MBBs with more than one reaction. In fact, the largest MBB in this core m-DAG encompasses 1039 reactions. In the plants’ cluster, i.e., cluster number 3, we again find that they share 631 MBBs from 1019 shared reactions. This implies that plants also share MBBs with more than one reaction, where in this case, the largest MBB in this core m-DAG includes 309 reactions. In addition, in cluster number 4 we obtained that only 2 reactions and the corresponding single MBBs are shared, namely, R03659 and R04773. Both are catalized by the enzyme 6.1.1.10 which corresponds to the methionine t-RNA ligase. The two common reactions correspond to a tRNA ligase. One of them is related to selenium metabolism and the other to methionine metabolism. Both compounds are fundamental to the metabolism of any living organism. This indicates that the organisms grouped in this cluster have distinct metabolisms.

In addition, we re-evaluated this classification by increasing the number of clusters to 5 and 6 clusters. In this case, we obtained that animals are further divided. Meanwhile, the organisms in cluster number 4, which consists of protists, fungi, algae, and nematodes, continue to be grouped. We refer to the pipeline available at *https://github.com/biocom-uib/MetaDag/tree/main* for a complete analysis of the hierarchical clustering results.

To further investigate the ability to classify m-DAGs at a deeper taxonomy level, we considered the Animals’ classification at the Phylum level. The corresponding heatmaps are displayed in Figure 4 on the right. We can observe that vertebrates are clearly separated from invertebrates. Also, within the invertebrates in the arthropods’ phylum, the class Insects are clearly differentiated from the others. Therefore, we can conclude that both m-DAGs similarities correctly classified the Eukaryotes at different taxonomy ranks. We refer to Files S8 to S13 in the Supplementary material for the heatmaps of the MSA and Munkres similarities at the Phylum level in the Plants, Fungi, and Protists kingdoms.

To end, we compared the agreement between the two proposed measures of m-DAG similarity. We calculated the Spearman and Pearson correlation coefficients between the values of MSA-similarity and Munkres-similarity obtained for each pair of m-DAGs. As a matter of fact, we obtained a value of 0.89 for the Spearman correlation coefficient and 0.91 for the Pearson correlation coefficient, indicating that both measures are nearly equivalent.

Figure 5 shows, in a box plot visualization, the similarity values of every pair of m-DAGs. As expected, the MSA-similarity measure obtained higher similarity values with a mean of 0,67 and a standard deviation of 0.18 while the Munkres-similarity measure obtained a mean of 0.55 with standard deviation of Hence, we conclude that both similarity measures almost equally classify the Eukaryotes within the different taxonomy groups.

**Figure 5:**
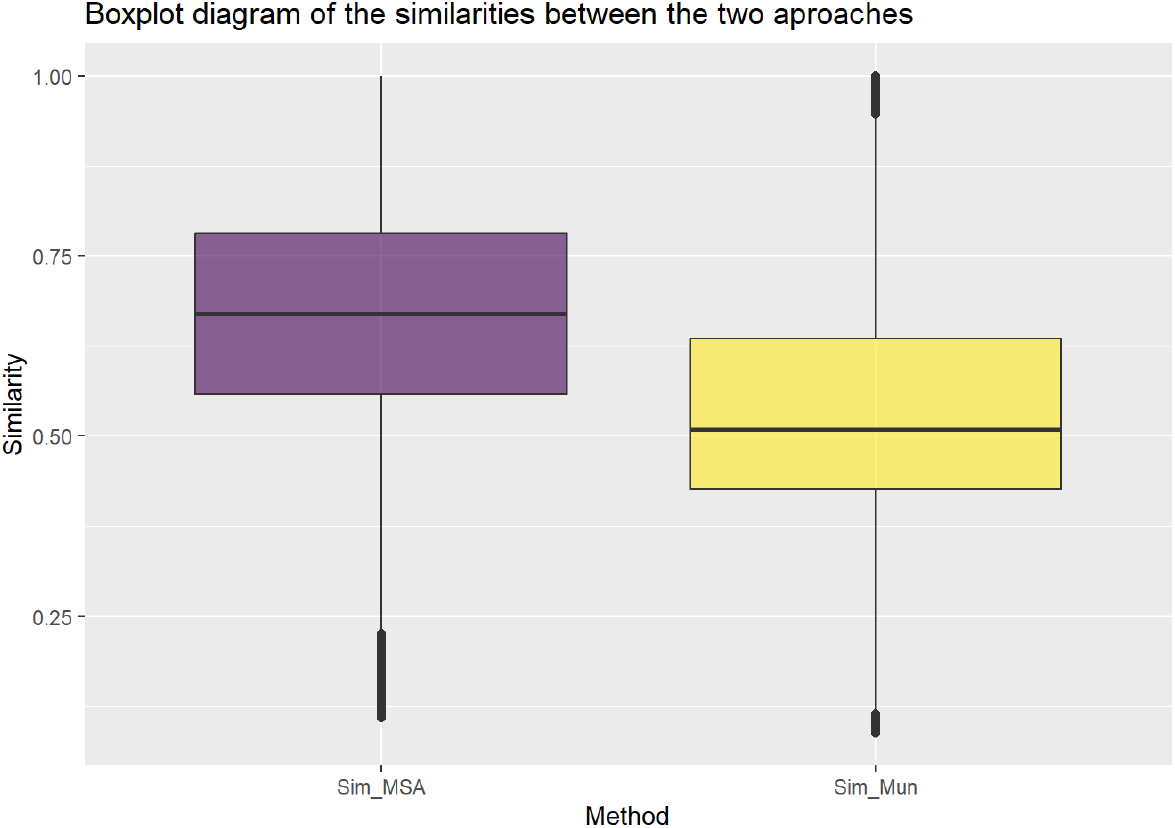
Boxplot of MSA-similarity values (left) and Munkres-similarity values (right) of the Eukariotes test.

## Conclusions

This paper introduces a robust implementation of the metabolic networks methodology, enabling the construction of metabolic Directed Acyclic Graphs (m-DAGs) to address diverse queries. These queries encompass analyzing specific pathways for individual organisms, exploring global metabolic networks of single organisms, studying pathways across all organisms in the KEGG database, investigating global metabolic networks for a list of organisms, examining synthetic metabolism involving various compounds, reactions, or enzymes, and facilitating comparisons between multiple experiments.

MetaDAG exhibits exceptional efficiency in rapidly computing reaction graphs and m-DAGs across a wide range of query types and data sources. Significantly, the integration with KEGG data empowers researchers to compare metabolisms associated with metagenomic and metatranscriptomic data, further enhancing the tool’s utility and versatility.

In addition, MetaDAG facilitates the construction of core and pan metabolisms from selected groups of experiments or organisms. This capability offers valuable insights into shared and distinct metabolic features, which contributes to understanding biological processes.

MetaDAG not only provides real-time interactive results on its user-friendly webpage but also facilitates further analysis and exploration through downloadable files. . In addition, we provide a comprehensive pipeline and guide to analyse the output results effectively. This resource equips researchers with the necessary tools and instructions to make the most of MetaDAG’s capabilities. We present the results of the Eukaryotes test, as well as MetaDAG’s performance and potential across a broad range of applications.

## Availability of data and materials

MetaDAG is available at: http://bioinfo.uib.es/metadag/home The supplementary material is available at: https://bioinfo.uib.es/MetaDAG_Supplementary_Material/

The pipeline to analyse MetaDAG results is available at: https://biocom-uib.github.io/MetaDag/.

